# Stress granule formation helps to mitigate neurodegeneration

**DOI:** 10.1101/2023.11.07.566060

**Authors:** M. Rebecca Glineburg, Evrim Yildirim, Nicolas Gomez, Xingli Li, Jaclyn Pak, Christopher Altheim, Jacob Waksmacki, Gerald McInerney, Sami J. Barmada, Peter K. Todd

## Abstract

Cellular stress pathways that inhibit translation initiation lead to transient formation of cytoplasmic RNA/protein complexes known as stress granules. Many of the proteins found within stress granules and the dynamics of stress granule formation and dissolution are implicated in neurodegenerative disease. Whether stress granule formation is protective or harmful in neurodegenerative conditions is not known. To address this, we took advantage of the alphavirus protein nsP3, which selectively binds dimers of the central stress granule nucleator protein G3BP (*rin* in *Drosophila*) and markedly reduces stress granule formation without directly impacting the protein translational inhibitory pathways that trigger stress granule formation. In *Drosophila* and rodent neurons, reducing stress granule formation with nsP3 had modest impacts on lifespan even in the setting of serial stress pathway induction. In contrast, reducing stress granule formation in models of ataxia, amyotrophic lateral sclerosis and frontotemporal dementia largely exacerbated disease phenotypes. These data support a model whereby stress granules mitigate, rather than promote, neurodegenerative cascades.

## Introduction

The integrated stress response (ISR) is an adaptive pathway by which any of four different serine/threonine kinases (GCN2, HRI, PERK, and PKR) respond to specific types of cellular stress and converge on a canonical translation initiation factor eIF2α. Phosphorylation of eIF2α inhibits ternary complex formation, which reduces global protein translation initiation without impacting elongation and thus creates large unassociated regions of mRNAs that can be recognized by cytoplasmic RNA binding proteins (Walter and Ron 2011; Starck et al. 2016). This process triggers the formation of stress granules (SGs), which are membraneless organelles that act as processing and sorting sites for mRNA to eventually allow for reinitiation or degradation following stress removal (Kedersha et al. 2000). The Ras-GAP SH3 domain binding protein (G3BP) is a key SG nucleator. It binds to both stalled 48S complexes via the 40s ribosome and interacts with single stranded RNAs exposed when translational initiation is blocked (Tourriere et al. 2003; Kedersha et al. 2016; Yang et al. 2020). These mRNAs are subsequently bound by other RNA binding proteins (RBPs) that associate with each other through low complexity domains (LCDs) and indirectly via RNA, creating a ribonucleoprotein complex (Kedersha et al. 1999; Kedersha et al. 2000; Buchan and Parker 2009). Overexpression of G3BP1 alone is enough to promote SG formation in the absence of stress, and inhibiting G3BP dephosphorylation prevents SG formation in response to ISR activation (Kedersha et al. 2016) (Tourriere et al. 2003; Yang et al. 2020).

Multiple converging points of evidence suggest that ISR activation and SG formation play prominent roles in neurodegenerative disease, particularly in amyotrophic lateral sclerosis (ALS) and frontotemporal dementia (FTD) (reviewed in (Baradaran-Heravi, Van Broeckhoven, and van der Zee 2020; Ling, Polymenidou, and Cleveland 2013; Wolozin and Ivanov 2019). One hypothesis is that SG formation accelerates RBP aggregation and that chronic ISR activation or SG protein-associated mutations impede SG disassembly, leading to formation of insoluble inclusions rather than dynamic liquid-liquid phase separation (LLPS) assemblies. In support of this hypothesis, mutations in genes encoding 11 SG proteins are associated with ALS or Alzheimer’s disease (Wolozin & Ivanova, 2019). These mutations tend to be gain-of-function mutations within LCDs that are associated with both translocation of the SG protein from the nucleus to the cytoplasm and protein aggregation (Barmada et al. 2010; Bilican et al. 2013; Serio et al. 2013; Johnson et al. 2008).

A key pathologic aggregate protein in ALS is TDP43, which accumulates in ubiquitin+ inclusions in >95% of ALS and approximately half of FTD cases (Arai et al. 2006).

TDP43 is an RBP that primarily resides in the nucleus, where it has key roles in mRNA processing. However, it moves to the cytoplasm in response to cellular stress pathway activation and is a component of some SGs (Fang et al. 2019; Colombrita et al. 2009; Ding et al. 2021). Over 40 mutations in the gene encoding TDP43 are linked with ALS and/or FTD. These mutations account for 2-5% of ALS/FTD cases and are overwhelmingly located within the C-terminal LCD. TDP43 mutants mis-localize to the cytoplasm, disrupt SG disassembly, and form persistent cytoplasmic protein aggregates (Ding et al. 2021; Barmada et al. 2010; Feneberg et al. 2020).

G_4_C_2_ hexanucleotide repeat expansions within the *C9orf72* gene are the most common mutations responsible for inherited ALS and FTD, accounting for up to 40% of familial disease (Renton et al. 2011; DeJesus-Hernandez et al. 2011). These expansions produce arginine-rich dipeptide repeats (DPRs) through a process known as repeat associated non-AUG (RAN) translation (Green et al. 2017; Lee et al. 2016; Mizielinska et al. 2014; Yamakawa et al. 2015; Flores et al. 2016). DPRs accelerate neurodegenerative pathology through a variety of mechanisms, one of which is associating with SGs and forming insoluble aggregates. Moreover, ISR activation enhances RAN translation and DPR production, and G_4_C_2_ repeat expression elicits ISR activation (Green et al. 2017; Lee et al. 2016; Sonobe et al. 2018; Cheng et al. 2018; Tseng et al. 2021; Westergard et al. 2019).

SG induction itself can also directly elicit neurodegeneration. ISR activation in the absence of a disease background recapitulates key ALS phenotypes including accumulation of TDP43 aggregates (Ratti et al. 2020; Fang et al. 2019; Zhang et al. 2019). Repeated formation of SGs in the absence of ISR activation, achieved by using optogenetics to artificially dimerize SG nucleator G3BP, also triggers formation of TDP43 aggregates and enhanced death in neurons (Zhang et al. 2019). These studies suggest that while mutant RBPs likely accelerate the rate of neurodegeneration, frequent ISR activation alone is sufficient to cause it.

If SG formation contributes to neurodegenerative disease, then inhibiting SG formation could be a viable strategy to prevent neurodegeneration. Indeed, a number of studies have used inhibitors of the ISR to suppress neurodegenerative phenotypes in mice and *Drosophila* (Fang et al. 2019; Das et al. 2015; Saxena, Cabuy, and Caroni 2009; Kim et al. 2014; Radford et al. 2015). However, blocking the ISR also prevents eIF2α triggered protein translational inhibition and activation of a stress response cascade that independently elicits cell death (Fang et al. 2019; Bordeleau et al. 2005; Dang et al. 2006; Kedersha et al. 2000), making it difficult to tease out the contributions of SGs specifically to neurodegeneration.

Viral proteins offer an intriguing alternative approach. Viruses have evolved methods to escape the ISR because its activation would otherwise prevent viral protein synthesis. Semliki Forest virus (SFV), escapes the ISR through generation of a nonstructural protein 3 (nsP3) which binds G3BP and prevents SG formation (Schulte et al. 2016; McInerney 2015; Panas et al. 2015; Panas, Ahola, and McInerney 2014; Panas et al. 2012; Gotte et al. 2019). Cells infected with WT SFV, but not with a mutant nsP3 SFV, were unable to form SGs in response to sodium arsenite (NaArs) and pateamine A, indicating nsP3 can inhibit both eIF2α dependent and independent SG formation (Panas et al. 2012; Dang et al. 2006). nsP3 binds NTF2-like domains of G3BP1 through its two FGDF domains, such that each FGDF domain can sequester one G3BP1 dimer to form a poly-complex with four G3BP molecules (Schulte et al. 2016). Binding to G3BP creates a network of nsP3:G3BP complexes that assist in viral replication, and promotes viral protein synthesis by recruiting 40S ribosomes to viral RNA (Gotte et al. 2019; Gotte et al. 2020; Schulte et al. 2016). Interestingly, a 31 amino acid fragment containing both FGDF domains (nsP3-31) is sufficient for binding G3BP1, and these domains are likewise necessary to inhibit SG formation by nsP3 (Panas, Ahola, and McInerney 2014; Panas et al. 2015; Panas et al. 2012).

Here, we took advantage of this nsP3-31 fragment to inhibit SG formation in ALS/FTD models and assess whether preventing SGs alleviated toxicity. While nsP3-31 effectively inhibited SG formation induced by a variety of mechanisms, it was detrimental in several neurodegenerative disease models, suggesting that SG formation may be beneficial at least initially in these contexts. These findings have important implications for future therapeutic development targeting these pathways in neurodegenerative disorders.

## Results

A GFP-nsP3-31 fusion protein was previously shown to bind tightly to G3BP (Panas et al. 2012). To determine whether this fusion protein inhibits canonical SG formation, we co-transfected either a WT EGFP-nsP3-31 (nsP3-WT) or a mutant EGFP-nsP3-31 (nsP3-mut) in which the two FGDF domains contain single phenylalanine (F) to alanine (A) mutations that prevent binding to G3BP (Panas et al. 2015; Panas et al. 2012). We then applied the dsRNA mimic poly(I:C), which activates the ISR through the dsRNA protein kinase PKR. Poly(I:C) robustly induced SGs when transfected into HEK293T cells (**Supplemental Figure 1**). NsP3-WT, but not nsP3-mut, inhibited poly(I:C) induced SGs (**Figure 1A-B**).

**Figure 1.**
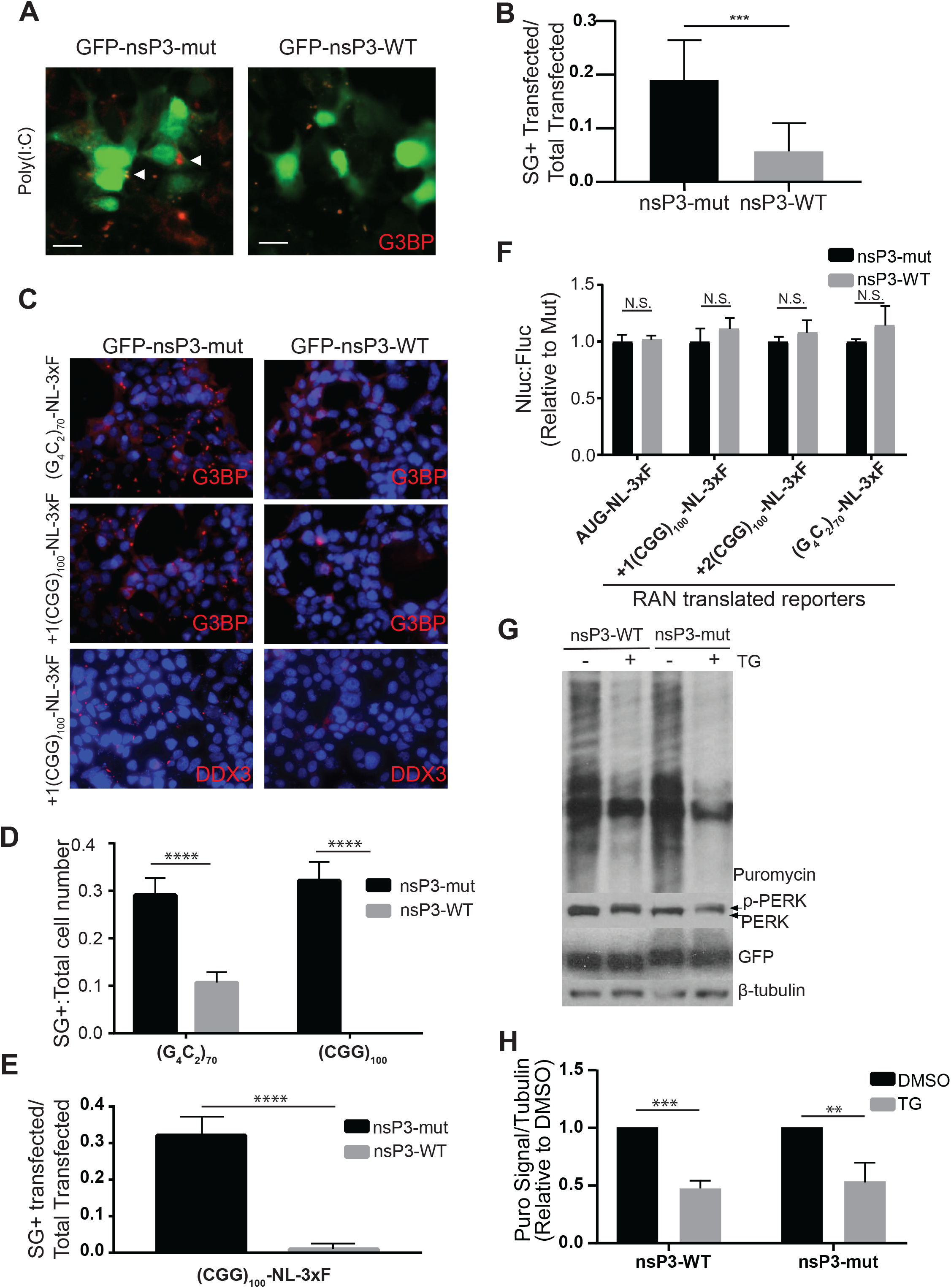
nsP3 inhibits SG formation caused by GC-rich repeats, but has no effect on translation. A) HEK 293T cells co-transfected with WT or mut nsP3 and poly(I:C). B) Quantification of G3BP SG+ cells in (A). Bars represent proportion +/- 95% C.I. mut nsP3 n= 117, WT nsP3 n=124. C) HEK 293T cells co-transfected with WT or mut nsP3 and the indicated Nluc-3xF reporters. G3BP and DDX3 are used as SG markers. D) Quantification of G3BP SG+ cells in (C). Bars represent proportion +/- 95% C.I.s (G4C2)70 + mut nsP3 n=622, (G4C2)70 + WT nsP3 n=731, (CGG)100 + mut nsP3 n=538, (CGG)100 + WT nsP3 n=685. E) Quantification of DDX3 SG+ cells in (C). Bars represent proportion +/- 95% C.I. (CGG)100 + mut nsP3 n=324, (CGG)100 + WT nsP3 n=314. (B, D- E) Two tailed fisher’s exact test with Bonferonni correction for multiple comparisons (D), ***p<0.001, ****p<0.0001. F) Nanoluciferase expression relative to FireFly luciferase (FFLuc) of indicated Nanoluciferase (Nluc)-3xF reporters co-transfected with EGFP-nsp3 WT or EGFP-nsP3 mut. Bars represent mean +/-SEM. Two-way ANOVA and Holm-Sidak unpaired t-tests, n=6. G) Western blot and SUnSET Assay (puromycin) of HEK 293Ts transfected with EGFP-WT nsP3 or EGFP-mutant nsP3 and treated with either DMSO (control) or Thapsigargin (TG). B-tubulin= loading control, p- PERK= control for TG ISR induction. H) Quantification of SUnSET assay in (G). Bars represent mean +/- standard deviation, Welch’s t-tests, n=3, *p<0.05, **p<0.01.

Next, we assessed whether nsP3-WT could inhibit SGs induced by a neurodegenerative trigger. We previously showed that Fragile-X-associated tremor ataxia syndrome (FXTAS) CGG-repeat and ALS/FTD associated G_4_C_2_-repeat containing RAN reporters activate the ISR and elicit eIF2α-dependent SGs (Green et al., 2017). Co-transfecting nsP3-WT with either FXTAS associated (CGG)_100_ repeats or ALS/FTD associated (G_4_C_2_)_70_ repeats significantly inhibited RAN reporter-induced SGs compared to nsP3-mut (**Figure 1C-E**). Despite inhibiting SG formation, the presence of nsP3-WT had no effect on canonical translation of AUG-nanoluciferase or firefly luciferase reporters or RAN translation of both CGG and G_4_C_2_ repeat containing reporters (**Figure 1F**). When cells were stressed with the ER calcium pump inhibitor, thapsigargin (TG), which activates PERK, neither nsP3-WT or nsP3-mut affected PERK activation or global translational inhibition (**Figure1G-H**). Together, these data suggest that nsP3 can be used to selectively inhibit SG formation without altering ISR-induced translational suppression.

To better understand the role of SGs in mammalian neurodegeneration, we investigated whether nsP3 could influence SG formation in rat cortical neurons. Consistent with prior studies, we confirmed the formation of SGs in rodent primary cortical neurons in response to NaArs using a whole-cell measure of granularity, as determined by the single-cell coefficient of variability (CV) (Khalfallah et al. 2018; Sharkey et al. 2018)(**Figure 2A**). In comparison to inactive nsP3-mut, nsP3-WT significantly reduced the appearance of SGs in transfected neurons (**Figure 2B**).

**Figure 2.**
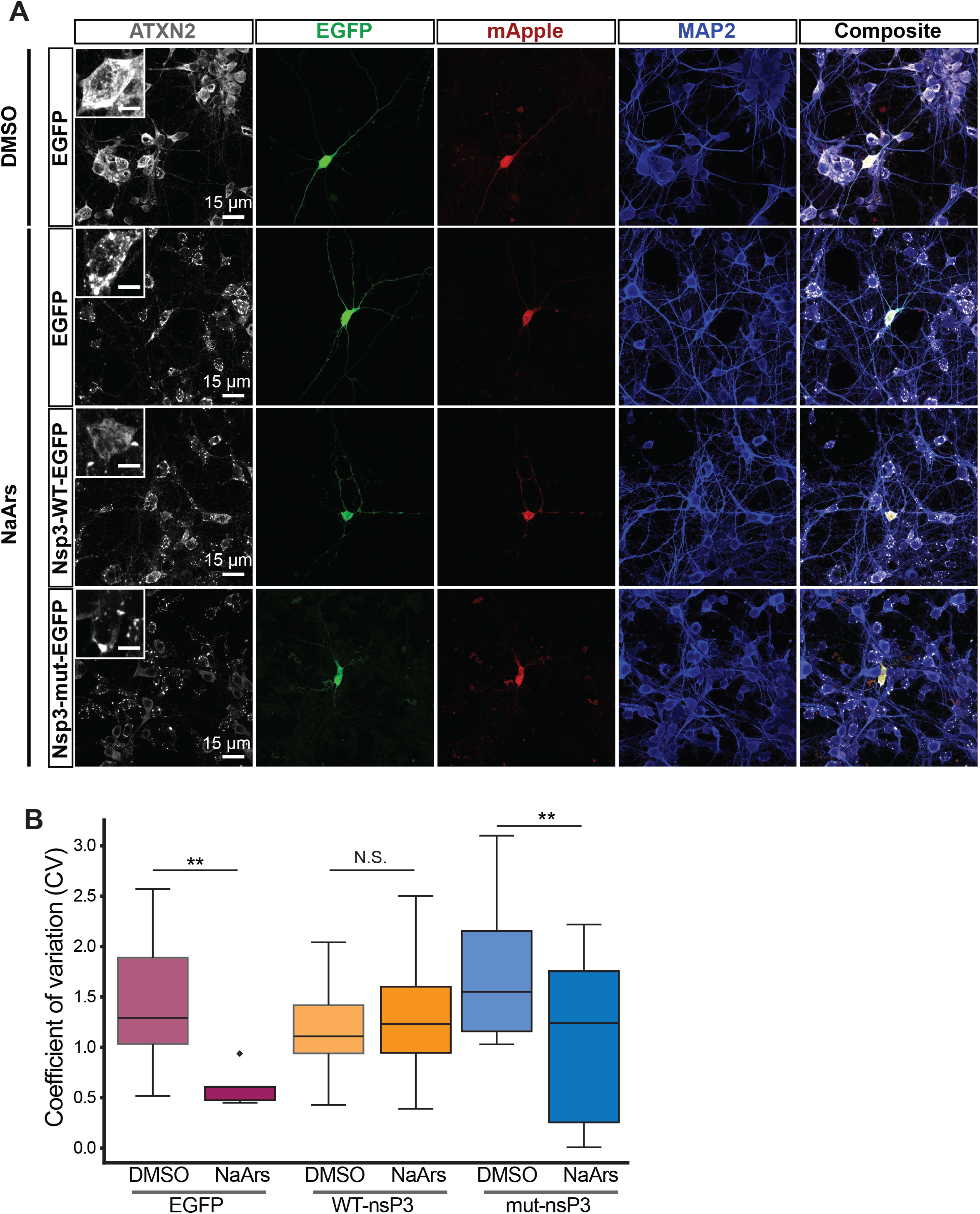
nsP3 reduces SG formation in neurons. A) Primary rat cortical neurons treated with 250mM NaArs for 30min prior to immunostaining for the SG marker ATXN2. B) Quantification of coefficient of variation (CV) of granular ATXN2 staining in (A). *p=0.021, **p=0.010, ns: p>0.05; Welch’s unpaired t-test. A. N=14-19 cells/condition, pooled from 3 replicates.

We next determined whether nsP3-WT expression could enhance survival in two established disease models: a TDP43 overexpression ALS/FTD model (TDP43- mApple), and a neuronal model of FXTAS ((CGG)_100_-EGFP) (Barmada et al. 2015; Linsalata et al. 2019). As expected, overexpressing TDP43-mApple significantly increased the risk of death in comparison to mApple alone (**Figure 3A**). Co-expression of nsP3-WT failed to rescue toxicity arising from TDP43-mApple, however, and instead increased the risk of death further. Similar results were observed with nsP3-mut overexpression. In addition, neither nsP3-WT nor nsP3-mut prevented neurodegeneration in our FXTAS rat primary neuron model (**Figure 3B**). Here, overexpression of (CGG)_100_-EGFP resulted in toxicity that was unaffected, and even somewhat exacerbated, by co-expression of nsP3-WT or nsP3-mut co-expression. Further, expression of nsP3-WT and nsP3-mut displayed subtle increases in the risk of death compared to EGFP control, suggesting intrinsic toxicity associated with the C-terminal 31 residues of nsP3 regardless of its ability to bind G3BP (**Figure 3A-B**). These data suggest nsP3-31-WT, despite reducing SG formation in neurons (**Figure 2**) fails to mitigate neurodegeneration associated with TDP43 or CGG repeat expression.

**Figure 3.**
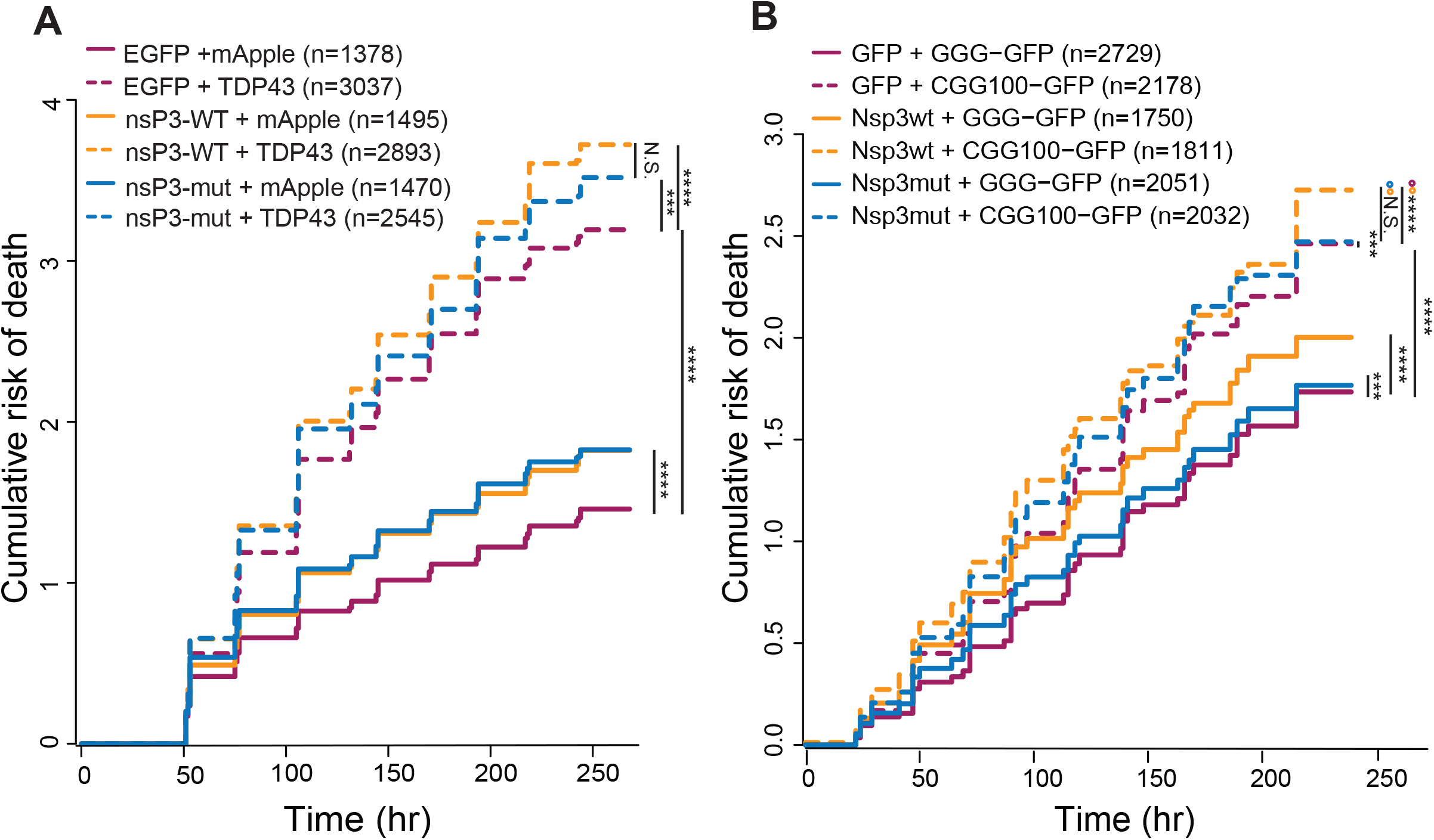
nsP3 does not enhance survival in primary neurons expressing TDP43 or trinucleotide repeats. A-B) Automated microscopy and survival analysis of neurons expressing A) TDP43-mApple and nsP3-WT, nsP3-mut, or mApple EGFP control, or B) (CGG)100-GFP and nsP3-WT, nsP3-mut, or mApple EGFP control. Data inA pooled from 3 biological replicates, each with 8 technical replicates/condition. Data in B are combined from 4 biological replicates, each with 8 technical replicates/condition. Cox proportional hazards analysis, ***p<0.001, ****p<0.0001.

To determine if nsP3 has any effects in an *in vivo* setting, we turned to a *Drosophila* model system. *Drosophila* homologue rin NTF2-like domains are only 60.6% identical to human G3BP1 and share identical or similar homology for 8 out of the 10 known nsP3 interacting sites (**Supplemental Figure 2A**) (Schulte et al. 2016); however nsP3 has also been shown to bind *Aedes albopictus* (mosquito) rin (Nowee et al. 2021; Fros et al. 2015), and *Drosophila* rin NTF2 domains are 93% identical to mosquito rin, including identical amino acid identity at all 10 nsP3 interacting sites (Supplemental Figure 2A). Thus, we reasoned that nsP3 could bind rin and also block SGs in *Drosophila*. We generated UAS-mApple-nsP3-WT (nsP3-WT) and UAS-mApple-nsP3-mut (nsP3-mut) *Drosophila* lines and compared overall expression levels to an UAS-mApple line (control). nsP3-WT lines had lower mRNA levels compared to control and nsP3-mut lines (**Supplemental Figure 2B**); however, mApple protein levels of both control and nsP3-WT were consistently higher than nsP3-mut (**Supplemental Figure 2C-E**). Importantly, when we coexpressed nsP3-WT with Rin-GFP, we observed no effect on total Rin-GFP levels compared to coexpression with nsP3-mut or mApple control, suggesting any differences observed between these three fly lines is not due to expression levels of Rin protein (**Supplemental Figure 2E**).

To determine if nsP3-WT interacts with rin and inhibits SG formation in flies, we dissected brains from nsP3-WT, and nsP3-mut 3^rd^ instar larvae expressing an endogenously GFP-tagged rin (Sarov et al. 2016) and exposed them to specific stress conditions. Both Actin-Gal4 expressing control and nsP3-mut lines showed strong SG formation in the central brain after heat shock or treatment with NaArs or thapsigargin (**Figure 4A and Supplemental Figure 3A-B**). In contrast, Actin-Gal4/nsP3-WT brains had no detectable SGs under any of these conditions. SGs were visible in cells in the optic lobes, but were rarely seen in the ventral nerve cord for all genotypes under all conditions (data not shown).

**Figure 4.**
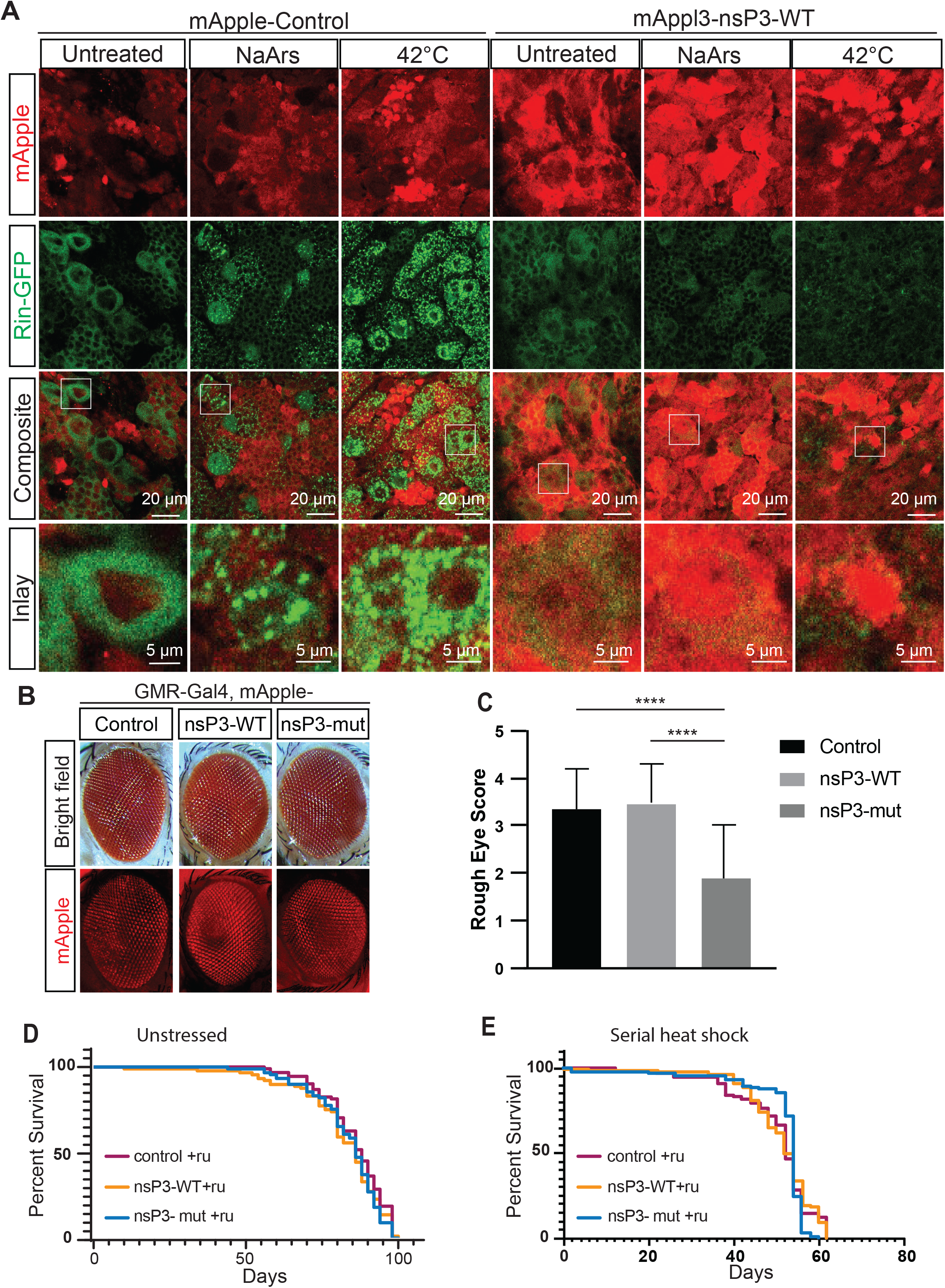
Impact of nsP3 expression in Drosophila. A) Representative images of 3rd instar larval brains expressing rin-sfGFP, and Actin-Gal4, UAS-mApple-control (left) or Actin-Gal4, UAS-mApple-nsP3 WT (right) untreated, 2 hrs in 500uM NaArs, or 1 hr at 42C. B) Representative images of eyes from flies coexpressing GMR-Gal4 with mApple- control,mApple-nsP3-WT, or mApple-nsP3-mut. Top) Bright field, Bottom) mApple expression. C) Quantification of rough eye phenotypes of males from B (Control n= 23, nsP3-WT n=33, nsP3-mut n=26). One-way ANOVA and Tukey’s multiple comparisons test. ****p<0.0001. D) Survival assay of Tub5-GS, mApple control, nsP3-WT, or nsP3-mut flies exposed to RU-486 at 24C. (control n= 92, nsP3-WT n= 90, nsP3-mut n= 89). Log-rank Mantel-Cox test. E) Survival assay Tub5-GS, mApple control, nsP3-WT, or nsP3-mut flies exposed to RU-486 and serially heat shocked (incubated at 24C then heat shocked for 2 hrs at 36C every 48 hrs). (control n= 90, nsP3-WT n= 89, nsP3-mut n= 88). D-E: Log-rank Mantel-Cox test, no significance observed between genotypes.

Expression of a variety of factors associated with neurodegeneration in the fly eye can elicit a rough-eye phenotype associated with ommatidial degeneration. GMR-Gal4 expression of nsP3-WT in isolation did not induce eye degeneration compared to control or nsP3-mut (**Figure 4B-C**). However, both the control and nsP3-WT expressing fly eyes scored significantly higher than nsP3-mut lines, which we attribute to differences in protein production (**Figure 4B-C, Supplemental Figure 2C-D, Supplemental Figure 3C**). As we were unable to balance our protein expression in these models, we used both control and nsP3-mut lines when comparing effects of nsP3. To assess the impact of SG formation inhibition on longevity, we utilized the Tub5-Geneswitch system to activate nsP3-WT, nsP3-mut, or mApple control expression ubiquitously post eclosion. Under non-stressed conditions at 24C, nsP3-WT alone had no impact on longevity in adult flies in comparison to nsP3-mut or control flies (**Figure 4D**). Similarly, Tub5-GS/nsP3-WT lines which were not treated with the gene-activating agent RU-486 lived similar lifespans compared to lines where Ru-486 activated nsP3- WT expression (**Supplemental Figure 3D compared to Figure 4D**). nsP3-WT expression also had no effect on eclosion proportions when expression was driven ubiquitously during development using an Actin-Gal4 driver (**Supplemental Figure 3E**). Taken together, these data suggest that nsP3 expression under non-stress conditions is not harmful.

To assess whether nsP3 might affect survival under conditions of serial stress exposure, we exposed flies expressing nsP3-WT, nsP3-mut, or mApple in neurons to serial heat shock (2 hours at 36 degrees every 2 days). This led to a significant decrement in survival in all fly lines; however, this effect was independent of genotype (**Figure 4E compared to 4D**). Thus, inhibition of SG formation even in the setting of serial stress response activation has no impact on survival.

We next investigated whether drivers of neurodegeneration associated with altered SG dynamics might respond to nsP3 expression. TDP43 aggregation in particular is a common feature in both ALS and FTD, and TDP43 itself is a SG protein. Moreover, pathogenic TDP43 cytoplasmic aggregation in ALS may be driven in part by SG formation (Ratti et al. 2020; Mann et al. 2019; Liu-Yesucevitz et al. 2010; Fang et al. 2019; Zhang et al. 2019). To investigate this in our model system, we co-expressed mApple control, nsP3-WT, or nsp3-mut with TDP43-EGFP in 3^rd^ instar larval brains using the neuroblast driver Asense-Gal4. This driver was chosen as previous attempts co-expressing nsP3-WT with TDP43-EGFP using the Actin-Gal4 driver was lethal prior to the 3^rd^ instar larval stage, and observations in rin-sfGFP flies indicated SG forming potential is abundant in neuroblasts (**Figure 4A**). Under non-stressed conditions, TDP43-EGFP expression was largely diffuse with occasional aggregates (**Supplemental Figure 3F**). nsP3-WT did not alter TDP43-EGFP expression levels or aggregates compared to nsP3-mut or mApple control expressing lines under non-stressed conditions. Upon NaArs treatment, the diffuse TDP43-EGFP signal decreased and brighter puncta were observed (**Supplemental Figure 3G**). nsP3-WT had no effect on number or size of TDP43-EGFP foci in brains compared to either the nsP3-mut or mApple control (**Supplemental Figure 3H-I**) This suggests nsP3-WT is unable to prevent stress-induced TDP43 aggregation.

We next asked whether nsP3-WT could alleviate neurodegenerative phenotypes in four *Drosophila* ALS models: two TDP43 models (overexpression of TDP43-EGFP or the ALS-associated TDP43 (M337V) mutant, which enhances TDP43 cytoplasmic mislocalization, disrupts SG dynamics and causes more rapid neurodegeneration in *Drosophila* (Ritson et al. 2010; Oh et al. 2015; Sreedharan et al. 2008; Feneberg et al. 2020)), and two *C9orf72* RAN translation models ((GGGGCC)_28_-GFP, and (GR)_100,_ both of which express glycine-arginine DPRs that form large GR aggregates that cause extensive neurodegeneration and impair SG dynamics (Feneberg et al. 2020; He et al. 2020; Mizielinska and Isaacs 2014; Lee et al. 2016)).

In GMR-Gal4 driven models that express TDP43-EGFP in the fly eye, nsP3-WT enhanced the rough eye phenotype compared to both control and nsP3-mut (**Figure 5A-B**). Similar effects were observed for both (GGGGCC)_28_-GFP, and TDP43(M337V) expressing lines **(Figure 5C-E and Supplemental Figure 4A-C**). nsP3-WT also reduced eye width and enhanced necrosis in (GGGGCC)_28_-GFP lines compared to control and nsP3-mut (**Figure 5F-G and Supplemental Figure 4D-E**) flies. However, nsP3-WT had no effect on the (GR)_100_ expressing line or a FXTAS RAN translation model (**Supplemental Figure 4F-G**). Together these results suggest that inhibiting SG formation and TDP43 aggregation is not beneficial, even in disease models where SG dynamics are known to be disrupted or where SG formation itself contributes to aberrant protein aggregation.

**Figure 5.**
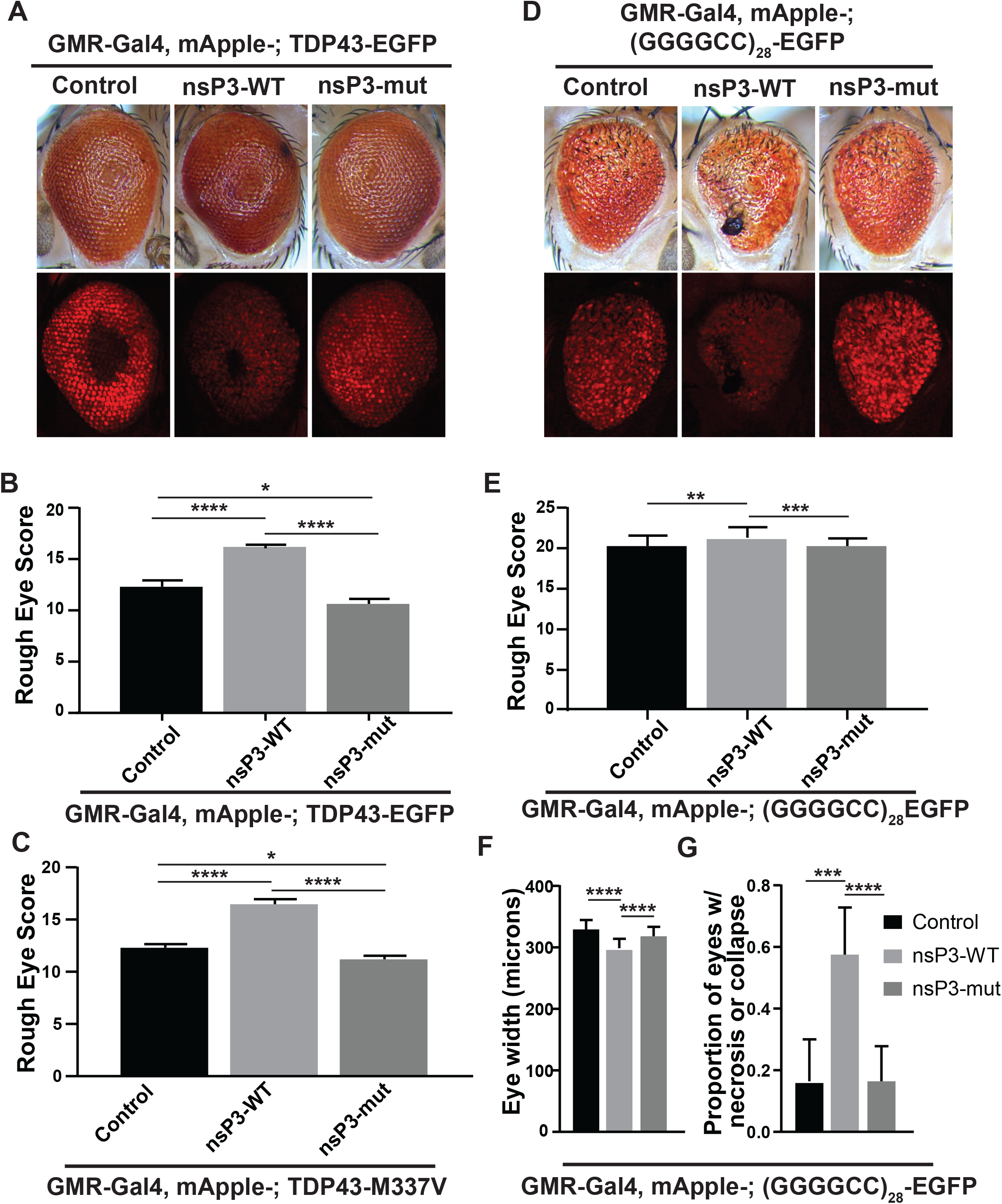
nsP3 enhances toxicity in ALS fly models. A) Representative images of male fly eyes expressing TDP43TDP43-EGFP under GMR-GAL4 expression with control, nsP3-WT, or nsP3-mut. B) Quantification of rough eye phenotypes of males from A (control n= 34, nsP3-WT n= 68, nsP3-mut n= 74). C) Quantification of rough eye phenotypes of males expressing TDP43-M337V under GMR-GAL4 expression with control, nsP3-WT, or nsP3-mut (control n= 33, nsP3-WT n= 33, nsP3-mut n= 52) ****p<0.0001. D) Representative images of male fly eyes expressing (GGGGCC)28-EGFP under GMR-GAL4 expression with control, nsP3-WT, or nsP3-mut. E) Quantification of rough eye phenotypes of males from D. (control n=44, nsP3-WT n=33, nsP3-Mut=61) . (F-G) Quantification of DF) eye width (control n= 23, nsP3-WT n= 23, nsP3-mut n= 21), and G) proportion of necrosis positive eyes (control n=44, nsP3-WT n=33, nsP3-Mut=61). B,C,E: One-way ANOVA and Tukey’s multiple comparisons test. F: Two-way ANOVA (see Supplemental Fig 5D) with Tukey’s multiple comparisons test. *p<0.05, **p,0.01, ***p<0.001, ****p<0.0001.G: Fischer’s exact test with Bonferonni correction for multiple comparisons. *p<0.01667, **p<0.01, ***p<0.001, ****p<0.0001.

We next investigated whether nsP3-WT could enhance longevity in stress and nonstress conditions in two ALS *Drosophila* models: a TDP43 overexpression fly, and *C9orf72* RAN translation model (GGGGCC)_28_-GFP) (He et al. 2020; Oh et al. 2015; Ritson et al. 2010). Both lines have significantly reduced survival relative to GFP control lines (**Supplemental Figure 5A**). nsP3-WT had no effect on longevity in flies expressing TDP43-EGFP compared to flies expressing nsP3-mut or mApple control under nonstress conditions (**Figure 6A *left***). Similarly, in this same TDP43 overexpression background, expressing nsP3-WT in the presence of serial heat shock exposure was also detrimental compared to nsP3-mut and mApple control (Figure 7A *right*). In the (GGGGCC)_28_-GFP background, nsP3-WT significantly reduced survival under non-stressed conditions (**Figure 6B *left***). Conversely, under serial heat shock exposure, nsP3-WT significantly improved survival of (GGGGCC)_28_-GFP flies, but not relative to nsP3-mut (**Figure 6B *right***). Whether this discrepancy is biologically relevant is unclear.

**Figure 6.**
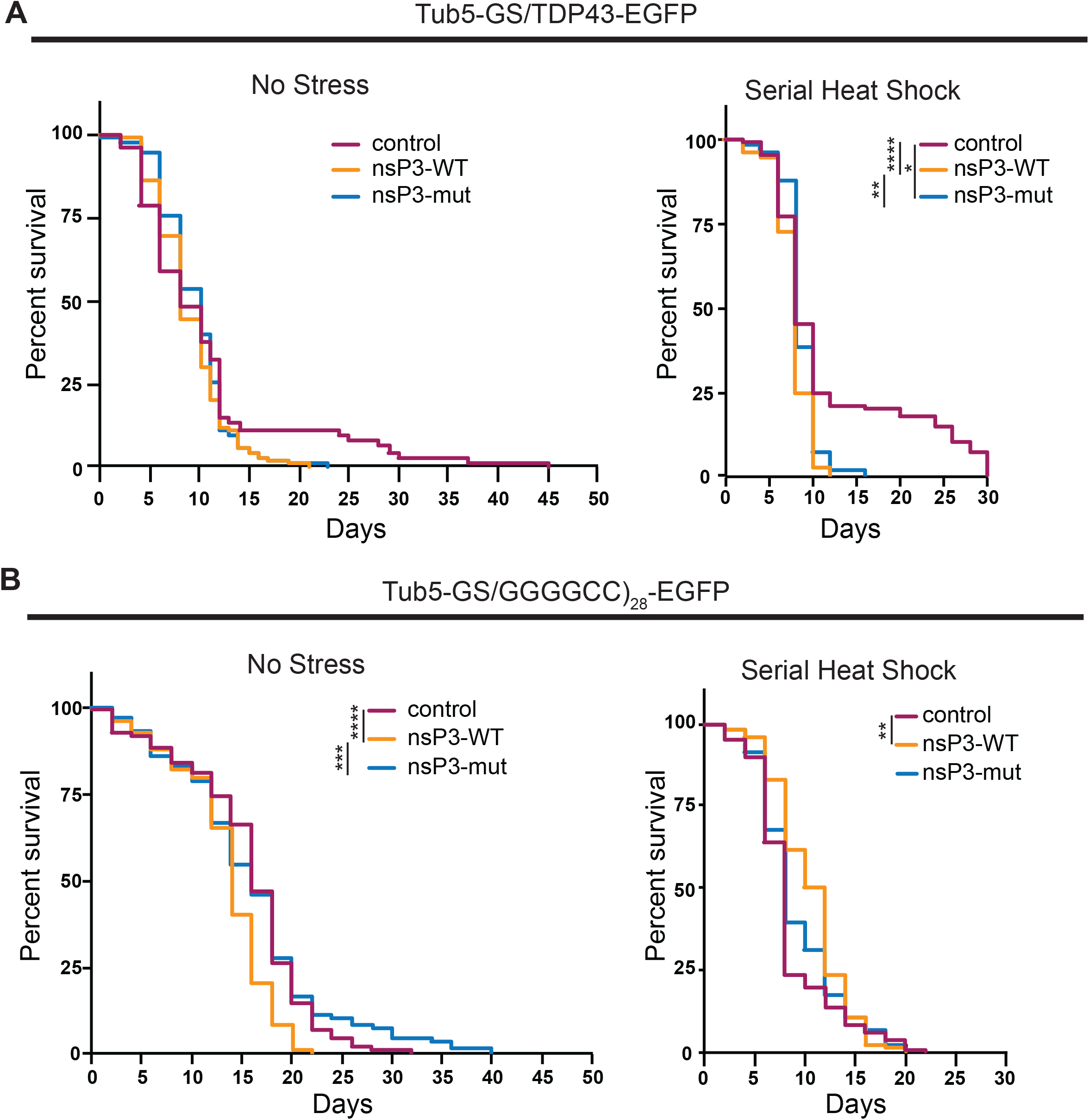
nsP3 does not enhance survival in ALS fly models. A) Survival curves of Tub5-GS/TDP43- EGFP coexpressed with indicated genotypes either at (left) constant 29C (control n= 75, nsP3-WT n=111, nsP3-mut n=112), or (right) serial heat shock (2 hrs every 48 hours at 36C) (control n= 81, nsP3- WT n=85, nsP3-mut n=76). B) Survival curves of Tub5-GS/(GGGGCC)28-EGFP coexpressed with indicated genotypes either at (left) constant 29C (control n= 87, nsP3-WT n=84, ns3P3-mut n=109), or (right) serial heat shock (2hrs every 48 hours at 36C) (control n= 111, nsP3-WT n=106, nsP3-mut n=106). Log-rank Mantel-Cox test with Bonferroni corrections for multiple comparisons. *p<0.0125, **p<0.01, ***p<0.001, ****p<0.0001.

FXTAS-associated CGG repeats have also been implicated in ISR activation, and expression of CGG repeats in flies produces polyglycine aggregates {Green, 2017 #196}{Todd, 2013 #213}. As with other models, nsP3-WT had no effect on survival of the FXTAS model fly expressing (CGG)_90_-EGFP (**Supplemental Figure 5B**). Together, these findings suggest that toxicity in these models occurs independent of SG formation.

## Discussion

Stress granule formation and dynamics accelerate multiple neurodegenerative processes, and several proteins and transcripts contained within SGs are implicated in neurodegenerative disease (Wolozin and Ivanov 2019; Zhang et al. 2019; Ratti et al. 2020; Fang et al. 2019). However, SGs also play neuroprotective roles in response to acute stressors (Koppenol et al. 2023; Shelkovnikova et al. 2013; McGurk et al. 2018; Mann et al. 2019; Gasset-Rosa et al. 2019). Here we took advantage of a viral peptide, nsP3, to specifically inhibit SG formation in a variety of model systems without impacting ISR pathways or global protein translation, which has been a potential confounder in prior studies (Fang et al. 2019; Das et al. 2015; Saxena, Cabuy, and Caroni 2009; Kim et al. 2014; Radford et al. 2015). Although nsP3-WT consistently inhibited SG formation, it was unable to alleviate neurodegenerative pathology in ALS/FTD and FXTAS models. Instead, inhibiting SG formation largely had no effect or enhanced toxicity, suggesting that SGs may play important protective roles in neurodegenerative cascades.

Many neurodegeneration-associated proteins (e.g. TDP43 and Tau) can associate with SG markers in patient tissues (Dormann et al. 2010; Liu-Yesucevitz et al. 2010; Mackenzie et al. 2017; McGurk et al. 2014; Umoh et al. 2018). Given their LLPS properties and the presence of SG markers in pathological inclusions, SG formation is thought to promote inclusion formation in multiple neurodegenerative diseases (Fernandes et al. 2020; Vanderweyde et al. 2012). Indeed, promoting SG formation in the absence of stress through use of optogenetic tools that drive G3BP dimerization is sufficient to trigger formation of inclusions (Zhang et al. 2019). However, we observed TDP43 to readily aggregate during acute stress even when SG formation was inhibited (**Supplemental Figure 3G**). This is consistent with other studies showing TDP43 and Tau form aggregates independent of SG formation (Mann et al. 2019; Gasset-Rosa et al. 2019; Fernandes et al. 2020). How then might SGs mediate neurodegenerative cascades?

Formation of TDP43 and polyglutamine aggregates is typically protective rather than detrimental to survival of cultured neurons (Arrasate et al. 2004; Barmada et al. 2010), and sequestration of mutant FUS to stress granules reduces levels of toxic species in the cytoplasm (Shelkovnikova et al. 2013). Thus, SGs may serve as repositories to buffer accumulation of toxic oligomeric species which drive neurodegeneration. In this scenario, selective inhibition of SG formation would be expected to enhance accumulation of these proteins in their most toxic forms, which would enhance neuronal death as we observe here. Consistent with this model, the low complexity domains that allow SG proteins to interact have recently been shown to promote non-fibrillar elastic solids over time, while fibrillar-like structures typically form outside of these granules (**PREPRINT** Alshareedah et al. 2023).

Both the unfolded protein response and heat shock response are activated in multiple neurodegenerative contexts—typically in association with elevated levels of p-eIF2α and reductions in global protein synthesis (Colombrita et al. 2009; Hoozemans et al. 2007; Chang et al. 2002; Ahmed et al. 2023; Ilieva et al. 2007). Yet, SGs are not typically observed within neurodegenerative tissue samples at autopsy (Colombrita et al. 2009). While there may be technical reasons for this (Sanchez et al. 2021), perhaps the lack of SGs in pathologic samples reflects a failure in neurons to respond to chronic stress. In this scenario, pathologic aggregates of proteins like TDP43 or DPRs are remnants of earlier protective SG formation induced by cellular stress. In this study, preventing SG formation in response to initial stressors may accelerate induced cell death at earlier stages in the pathogenic process.

A beneficial role for SGs in neurodegenerative disease is supported by the fact that age is the most common risk factor for disease. Aging is associated with enhanced gene expression, dysregulated proteostasis, and increased oxidative damage, all of which exist as chronic activators of the integrated stress response (reviewed in Cao, Jin, and Liu 2020). Further, many proteins that are involved in neurodegeneration are normally required for proper SG dynamics. Loss of TDP43 inhibits robust SG assembly, while ALS/FTD causing *TIA1* mutations prevent SG disassembly (Khalfallah et al. 2018; Mackenzie et al. 2017). Lastly, multiple studies have shown that overexpression of mutant, aggregate-prone peptides in a variety of model systems can elicit the ISR within short timeframes (Green et al. 2017; Zhang et al. 2014; Tseng et al. 2008; Chafekar et al. 2008; Bellucci et al. 2011; Hoozemans et al. 2007; Hoozemans et al. 2012). Despite this accumulation of stressors and noted increases in p-eIF2α in neurons with age, SG formation efficiency appears to depreciate in older neurons and senescent cells (Khalfallah et al. 2018; Derisbourg, Hartman, and Denzel 2021; Moujaber et al. 2017). In this context, our findings suggest that inhibiting global translation in and of itself may be insufficient to combat aging-inducing insults, and SGs may play essential roles in promoting neuronal viability.

We and others have noted that cells pre-conditioned with acute or chronic stress exhibit impaired stress responses to exogenous stressors (e.g. sodium arsenite), due to elevated levels of the eIF2α dephosphorylase, GADD34 (Shelkovnikova et al. 2017; Kroschwald et al. 2015; Klein et al. 2022). If neurons within neurodegenerative tissue are chronically under stress, they may lose their ability to continually form and reform SGs in response to new stress events. Thus, neurodegeneration may not be the result of an overactive stress response, but rather, the inability to elicit a robust or consistent stress response over time (Cestra et al. 2017). This is supported by studies showing that loss of GADD34 enhances survival and delays neurodegeneration in various ALS model systems, and loss of PERK accelerates disease progression, while pharmacologically upregulating heat shock and unfolded protein response pathways significantly improves outcomes in ALS models (Ahmed et al. 2023; Saxena, Cabuy, and Caroni 2009; Wang, Popko, and Roos 2011, 2014; Das et al. 2015; Ghadge et al. 2020). A more recent study has also shown that ISR inhibition via eIF2B activators enhances disease progression in ALS models (Marlin et al. 2023). In the context of our own work, these studies support a predominantly protective role for the integrated stress response in neurodegeneration.

This study has some important limitations. First, it relies on overexpression disease models across relatively short time frames. Thus, these studies do not recapitulate the gradual accumulation of insults over months to years as likely occurs in neurodegeneration in humans. Second, our approach to prevent SG formation in these neurodegenerative models, while novel, has its own caveats: SG inhibition is incomplete in our model systems, and the nsP3-31 peptide itself appears in some contexts to have mild intrinsic toxicity, regardless of whether it can bind G3BP/rin or not (**Figure 3**). Thus, future studies using *in vivo* mammalian systems and orthogonal approaches will be needed to fully ascertain how generalizable these findings are to other contexts and settings.

Despite these caveats, our observations have potentially important implications for therapy development in neurodegenerative disease. If the ISR is an active participant in most neurodegenerative settings, then defining the distinction between SG formation and ISR induced translational repression and transcriptional activation within different neurodegenerative contexts will be important next steps in allowing us to target the correct nodes in these pathologic cascades. We propose that elucidating these nodes and defining their broader impact across the full time-course of neurodegenerative disease should both reveal novel therapeutic targets and help us avoid disruption of intrinsic neuroprotective cascades that may currently mitigate the severity and progression in these challenging conditions.

## Methods

### Plasmids and primers

pEGFP-nsP3-C-term-WT, and pEGFP-nsP3-C-term-F3A were previously generated (Panas et al. 2015; Panas et al. 2012; Schulte et al. 2016). pUASTattB and pUASz1.1 plasmids for making transgenic flies were generated using restriction enzyme cloning. pUASTattB plasmids were generated by inserting mApple and downstream C-term sequences between NotI and KpnI in pUASTattB parental vector (Bischof et al. 2007) (Table 1). pUASz1.1 plasmids were generated by inserting mApple and downstream C-term sequences between NotI and XbaI in pUASz1.1 (DGRC Stock 1433 ; https://dgrc.bio.indiana.edu//stock/1433 ; RRID:DGRC_1433). Primers were synthesized from IDT.

**Table 1:**
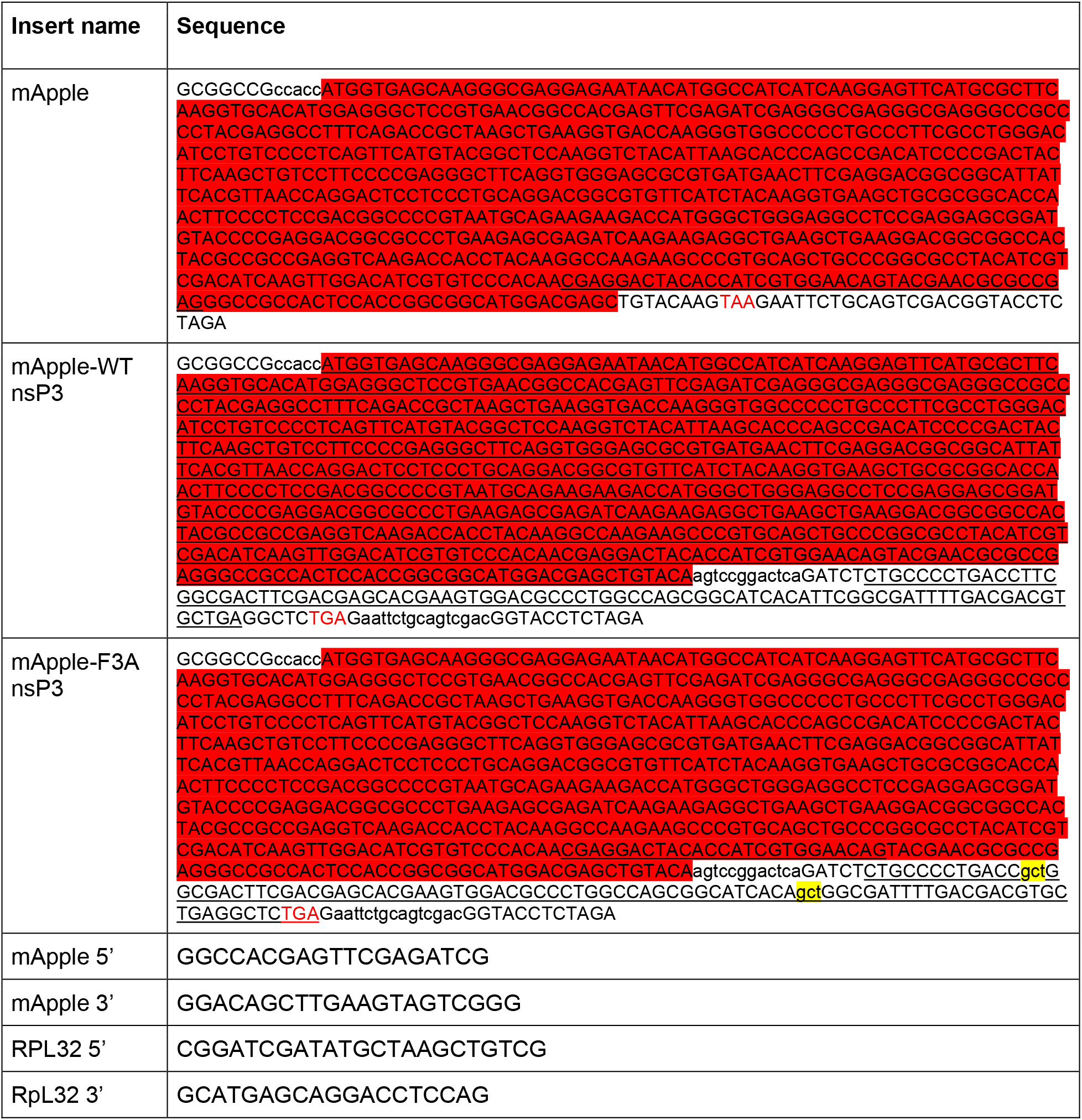
Plasmid and Primer Sequences used in this study.

**Table 2:** Transgenic fly lines used in this study.

| **Genotype** | **Figures** | **Source** |
| --- | --- | --- |
| ;UAS-mApple/CyO | Fig. 4B-E Supp. Fig. 3C-E | This study |
| ;UAS-mApple-nsP3-WT 2//CyO | Fig. 4B-E Supp. Fig. 3C-E | This study |
| ;UAS-mApple-nsP3-mut//CyO | Fig. 4B-E Supp. Fig. 3C-E, Fig. 5A, Fig. 6A, C, E Supp. Fig 4A, C | This study |
| ;UASz-mApple-nsP3-WT/CyO | Fig. 4A, Supp. Fig. 2B-D, Supp. Fig. 3B | This study |
| ;UASz-mApple-nsP3-mut//CyO | Fig. 4A, Supp. Fig. 2B-D, Supp. Fig. 3A-B | This study |
| UAS-TDP43-EGFP | Fig 5A, Fig 6A, C, Supp. 4A | This study |
| GMR-Gal4, UAS-CGG90-EGFP-BD/CyO | Supp. Fig. 4G Fig 6E, Supp Fig. 4C | (Jin et al. 2003) |
| w[1118];P{y[+t7.7] w[+mC]=UAS-poly-GR.PO-100}attP40 | Supp. Fig 4F Supp. Fig. 5B Fig 6E, Supp Fig. 4C | (Mizielinska and Isaacs 2014) |
| FlyFos028942(pRedFlp-Hgr)(rin[36711]:2XTY1-SGFP-V5-preTEV-BLRP-3XFLAG)dFRT | Supp. Fig. 4G Fig 6E, Supp Fig. 4C | VDRC 318907 (Sarov et al. 2016) |
| GMR-Gal4 |  | BDSC 8605 |
| Tub5-GeneSwitch |  | (Osterwalder et al. 2001) |

### Cell studies

Transfections were performed as previously described (Green et al., 2017). In brief, HEK293Ts were seeded at 1x10^5^ cells/ml and transfected with 3:1 FuGene HD 24hr later. For Nluc assays: a 1:1:10 ratio of Nluc reporter:FFluc reporter: nsP3 effector plasmid was used. For ICC, a 1:1 ratio of WT or mut nsP3 plasmid was co-transfected with either Poly(I:C) or Nluc reporter. 24 hrs post transfection, cells lysates were harvested for western blots, SUnSET assay, and Nanoluciferase assays as previously described or fixed in 4% PFA for immunocytochemistry (Green et al., 2017). The following antibodies were used for ICC: ms anti-DDX3 (1:250) (SCBT sc-81247), ms anti-G3BP1 (1:250) (BD Bioscience BDB611127), Goat anti-mouse Alexa Fluor 633 (1:500) (Invitrogen A-21146). The following antibodies were used at 1:1000 for HEK293T lysate western blots: ms anti-B-tubulin (DSHB E7), and ms anti-puromycin (Millipore MABE343), rb anti-GFP (Sigma 11814460001), and rb anti-PERK (CST 3192S). Blots were detected using HRP conjugated goat anti-mouse and goat anti-rabbit secondaries.

### Rough eye scoring & eye width measurements

Male flies containing the neurodegeneration causing transgene were crossed to GMR- Gal4 females at either 29C (UAS-(CGG)_90_-EGFP, TDP43-EGFP, TDP43-M337V), 25C ((GGGGCC)_28_-EGFP), or 18C (GR_100_-EGFP). One eye from each progeny was assayed for a rough eye phenotype or eye width measurement. Rough eye phenotype assays were performed according to previously published methods (Singh et al., 2021). Fly eye images were taken using a Leica M125 stereomicroscope and a Leica K3C equipped with LAS X manual z-stacking software. Eye widths were measured using LAS X software.

### Survival Assay

Fly longevity assays were performed according to previously published methods (Singh et al., 2021). Briefly, flies were separated by sex at 0-48 hrs post eclosion, and put on food containing RU-486. Every 48 hrs, dead flies were counted and surviving flies were moved to fresh food containing RU-486. Serial heat shock survival assays were performed similarly with the addition of exposing flies to 36C heat shock for 2 hrs every 48 hours.

### Larval qPCR and Western blots

10 3^rd^ instar wandering larvae for each genotype were collected and stored at -80C prior to lysis. Larvae were resuspended and homogenized in trizol according to manufacturer’s protocol (Invitrogen Trizol) with the following modifications. RNA was precipitated in isopropanol overnight at -20C. RNA was resuspended in 50uL H2O and column purified prior to cDNA synthesis.

500ng of RNA/sample was used to make cDNA using iScript cDNA synthesis kit according to manufacturer’s protocol (Bio-Rad). cDNA abundance was measured using 300nM of indicated primers and a iQ5 qPCR system (Bio-Rad).

For fly westerns, 5 3^rd^ instar wandering larvae/genotype were lysed in RIPA + miniComplete protease inhibitor cocktail, hand homogenized on ice 20x, and centrifuged at 12,000 xg and 4C for 5 minutes. Lysates were transferred to new tubes, homogenized through a 28G insulin syringe 8x, and boiled in 1xLB for 5 minutes.

Samples were run on 12% Bis-Tris gels, and transferred according to previously published protocols (Singh, et al., 2021). Membranes were blotted with rb anti-mCherry (ThermoFisher PA5-34974) 1:1000 (for mApple), ms anti-B-tubulin (DSHB E7) 1:1000, and ms anti-puromycin (Millipore MABE343), 1:1000, and detected using HRP conjugated secondaries Gt-anti-ms, and Gt-anti-rb 1:5000.

### Larval Brain Dissections, Drug treatments, and Fluorescence Imaging

Brains from 3^rd^ instar wandering larvae were dissected in 1xPBS, and transferred to Shields and Sang M3 insect medium, with 2% FBS and 2.5% fly extract (Gareau, C., Plos One, 2013). Dissected brains were incubated at 42C for 1 hr, or at 25C with either 500uM NaArs (2hrs), 10uM Thapsigargin (1.5 hrs), or untreated (2hrs). Brains were then washed in 1xPBS, fixed for 20 min in 4% PFA at RT, washed 3x for 5 min in 1x PBS, then incubated overnight at 4C in 1xPBS + 0,1% Tween with 1ug/ml DAPI, and mounted in Prolong Gold. For rin-sfGFP samples, Z-stacks containing 1uM sections were imaged on an Olympus FV100 using 40x oil objective, 2x zoom, and processed using ImageJ. 1 slice/brain is depicted in representative images. TDP43 samples were imaged on an Zeiss LSM 880 Confocal Microscope using 63x oil objective at 1.3 zoom, 1um pinhole, and processed using ImageJ as above. Foci over a specified threshold were counted as either small (5-15 pixels), medium (15-25 pixels), or large (25-35 pixels).

### Eclosion Assay

15 Actin/Cyo females and 5 mApple-x/mApple-x males were put on apple agar plates with wet yeast for 24 hrs. Embryos were collected at 24 hrs, counted under a microscope, and put on fresh 10% SY food. Number and genotype of eclosed flies were recorded 10-15 days later. The proportion of mApple expressing/total number eclosed was used to determine eclosion rates with each genotype, with an a priori expected ratio of 1:1 mApple expressing: Cyo, mApple not expressing.

### Primary neuron transfection

Primary mixed cortical neurons were dissected from embryonic day 19-20 Long-Evans rat pups and cultured at 0.6 x 10^6^ cells/mL in 96 well cell culture plates (TPP) coated with laminin (Corning) and D-polylysine (Millipore)(Archbold et al. 2018; Flores et al. 2019; Malik et al. 2018; Weskamp et al. 2019). At *in vitro* day (DIV) 3-4, neurons were transfected with 0.2 μg DNA and 0.5 μL Lipofectamine 2000 (ThermoFisher) per well, per the manufacturer’s protocol, with the exception that cells were incubated with Lipofectamine/DNA complexes for 20 min at 37**°**C before rinsing. Automated longitudinal fluorescence microscopy began 24h post-transfection for 10d, as previously described(Archbold et al. 2018; Flores et al. 2019; Malik et al. 2018; Weskamp et al. 2019; Arrasate et al. 2004). Briefly, images were acquired by an inverted Nikon Ti microscope equipped with a 20x objective lens, a PerfectFocus system, a Lambda 421 multichannel LED with 5 mm liquid light guide (Sutter), and either an Andor iXon3 897 EMCCD camera or Andor Zyla4.2 (+) sCMOS camera. All stage, shutter, and filter wheel movements were carried out by custom code written in publicly available software (μManager, ImageJ).

### Primary neuron imaging and puncta determination

Primary cortical neurons were isolated and plated onto glass coverslips, then transfected on DIV as described above. Forty-eight hours post-transfection, neurons were exposed to 250mM sodium arsenite or DMSO in PBS for 30 min at 37C, then washed twice with PBS before fixing with 4% paraformaldehyde in PBS for 10 min at RT, and permeabilizing with 0.1% Triton-X-100 in PBS for 20 min at RT. The cells were neutralized with 10 mM glycine in PBS at RT before blocking in PBS containing 0.1% Triton, 2% FCS and 3% BSA, for 1 h at RT. Detection was accomplished by incubating with anti-ATXN2 primary antibodies (diluted 1:500 in blocking solution, BD Biosciences 611378) overnight at 4°C. Samples were rinsed 3 times in PBS, incubated with Alexa 647 labeled donkey anti-rabbit secondary antibodies (1:250 in blocking solution; Invitrogen A32795TR) for 1 h at RT, then rinsed 3 more times in PBS containing a 1:1000 dilution of Hoechst 33258 nuclear dye (Invitrogen H3569) prior to confocal fluorescence imaging using a Nikon NSPARC confocal microscope. Sum intensity projections were created for each image, and regions of interest corresponding to individual neurons were drawn for GFP-positive soma. Within each region of interest, the coefficient of variability (CV) was calculated as the ratio of standard deviation of signal intensity to the mean signal intensity for the ATXN2 channel.

### Ethics statement

All vertebrate animal work was approved by the Committee on the Use and Care of Animals (UCUCA) at the University of Michigan, and the Louisiana State University Health Sciences Center at Shreveport’s Animal Care and Use Committee (ACUC). All experiments were performed in accordance with UCUCA and ACUC guidelines. Rats (*Rattus norvegicus*) used for primary neuron collection were housed singly in chambers equipped with environmental enrichment. Rats used for *in vivo* studies were housed with the dam until weaning at three weeks of age. Thereafter, they were housed in pairs by gender. All studies were designed to minimize animal use. Rats were cared for by the Unit for Laboratory Animal Medicine at the University of Michigan or veterinary specialists at Louisiana State University; all individuals were trained and approved in the care and long-term maintenance of rodent colonies, in accordance with the NIH-supported Guide for the Care and Use of Laboratory Animals. All personnel handling the rats and administering euthanasia were properly trained in accordance with the UM Policy for Education and Training of Animal Care and Use Personnel. Euthanasia was fully consistent with the recommendations of the Guidelines on Euthanasia of the American Veterinary Medical Association.

## Acknowledgements

Author Contributions

The project was initially conceived by PKT and GM and then designed and implemented by MRG with input from PKT, EY, NG and SJB. Experiments were conducted by MRG with assistance from EY, NG, JP, XL, CA, and JW. MRG wrote the manuscript with PKT and SJB and editorial input from all authors. Special thanks to Tahrim Chaudrey, Shannon Miller and Indranil Malik who assisted with survival assays.

