## Supplemental Figures for "Stress granule formation helps to mitigate neurodegeneration"

Supplemental Figure 1

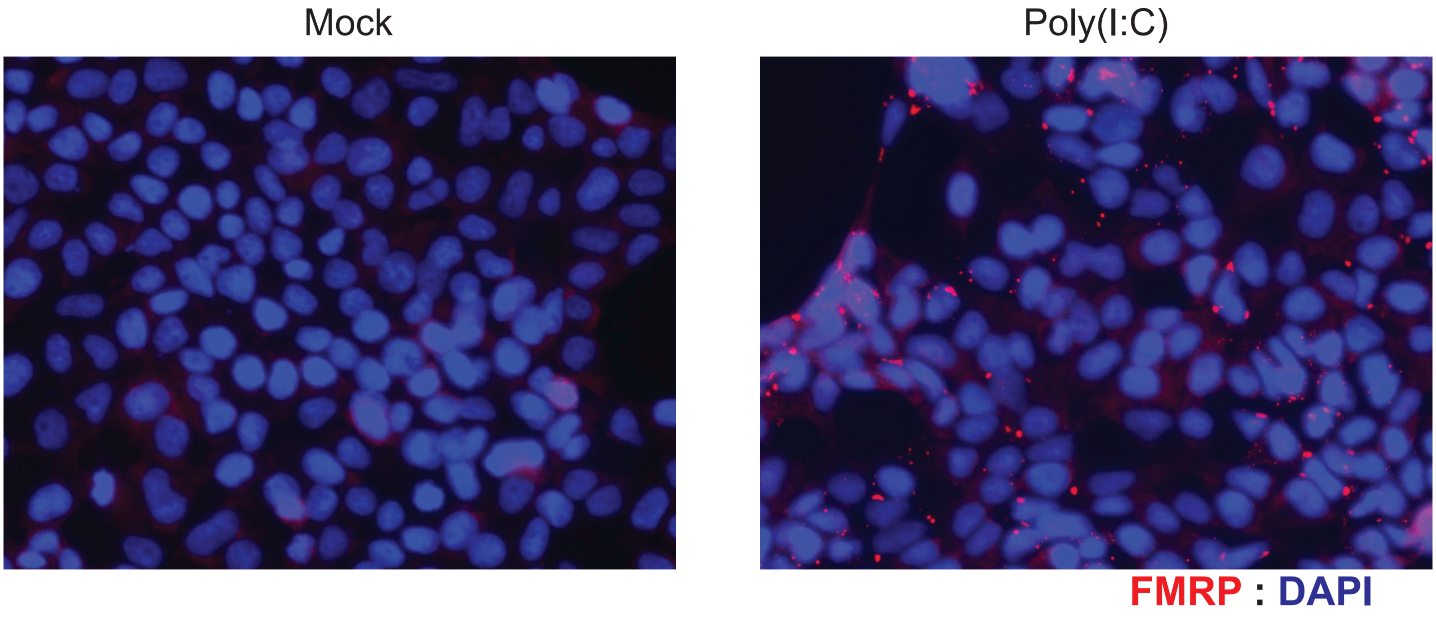


**Supplemental Figure 1. Poly(I:C) induces SGs.** Representative images of HEK 293Ts either mock or poly(I:C) transfected. FMRP= SG marker.

Supplemental Figure 2

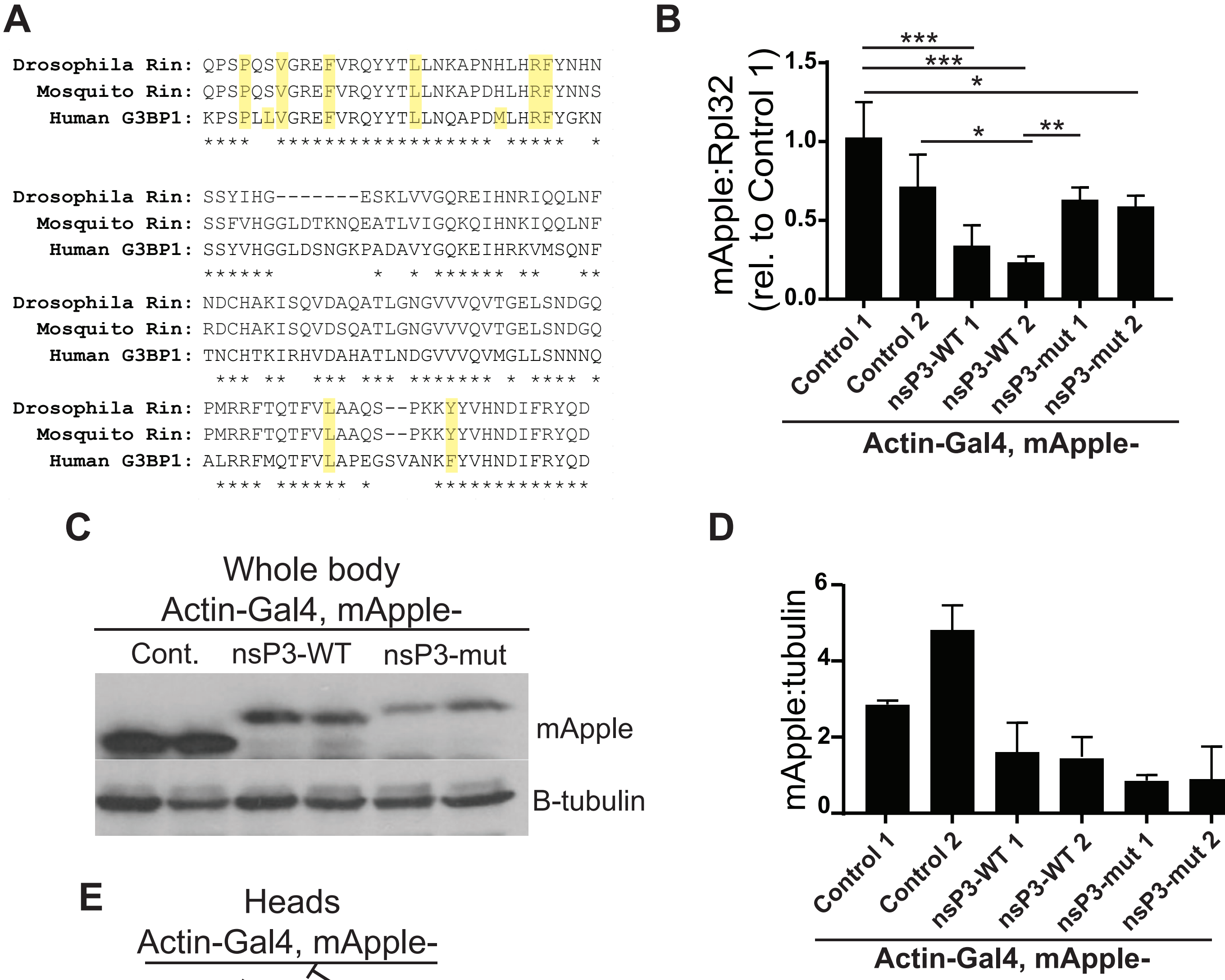


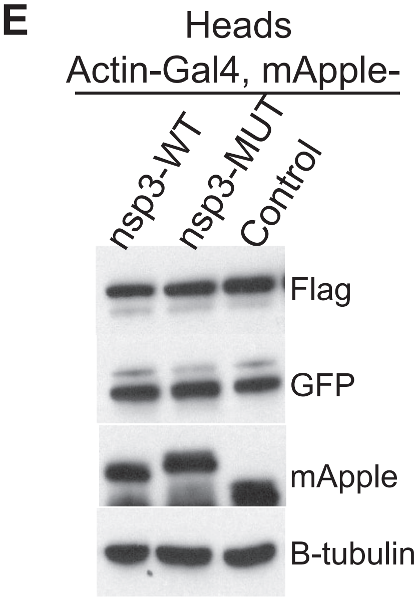


**Supplemental Figure 2: Characterization of nsP3 in Drosophila.** A) Schematic of Drosophila rin, Aedes albopictus rin and human G3BP1 NTF2-like domains. Asterisks indicate homology, yellow highlights denote nsP3 interacting regions. B) mRNA levels in Actin-Gal4, mApple-control, nsP3-WT, or nsP3-mut expressing 3rd instar larvae relative to Rpl32. Bars represent mean +/- standard deviation. One-way ANOVA with Tukey's multiple comparisons test, n=3 *p<0.0125, **p<0.01. C) Representative western blot of lysates from Actin-Gal4, mApple-control, nsP3-WT, or nsP3-mut 3rd instar larvae. B-tubulin= loading control. D) Quantification of western blot in (C). Bars represent mean+/- standard deviation, n=2.E) Representative western blot of lysates from Actin-Gal4, mApple-control, nsP3-WT, or nsP3-mut adult heads. B-tubulin= loading control. Flag and GFP recognize rin-sfGFP

Supplemental Figure 3

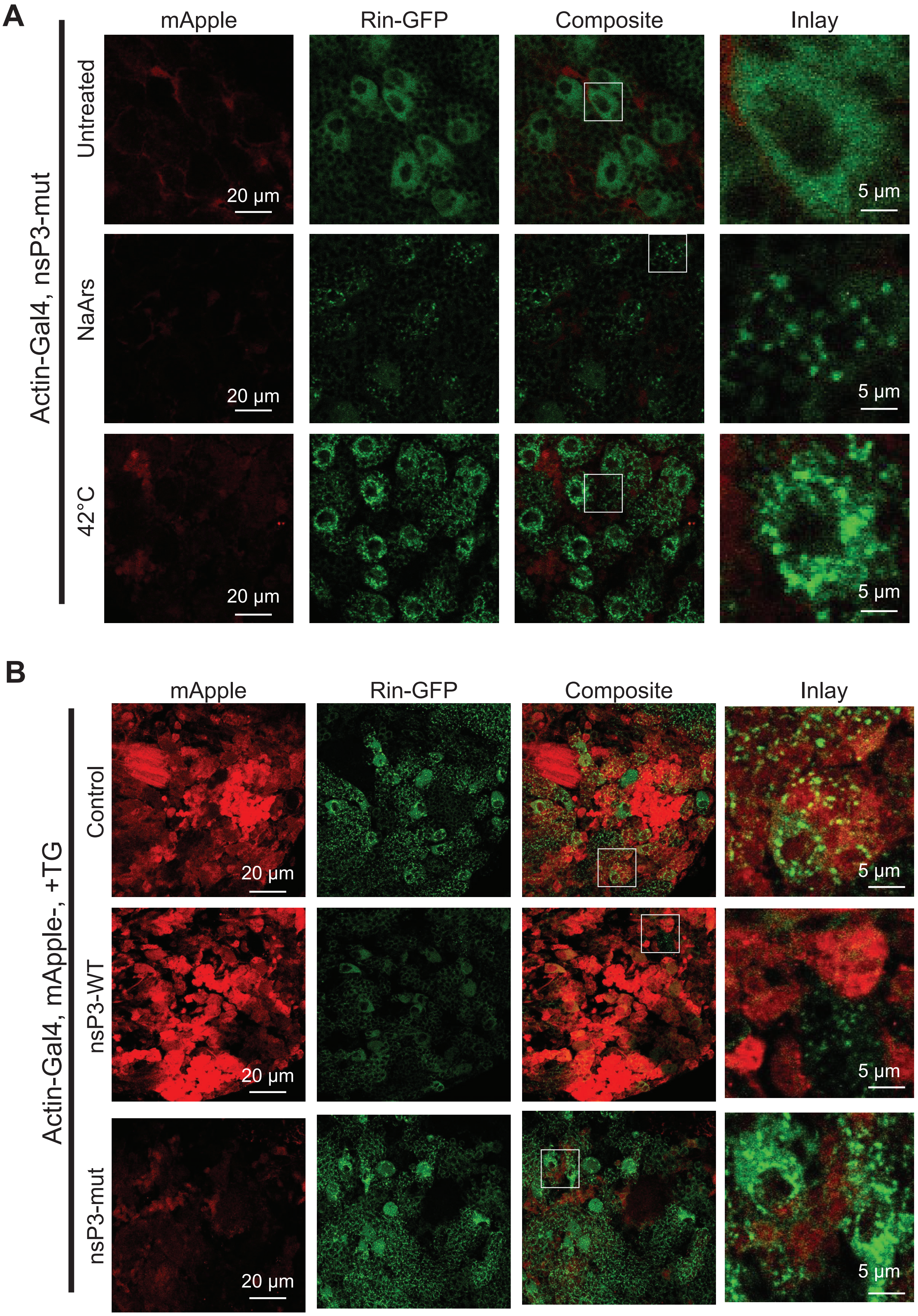

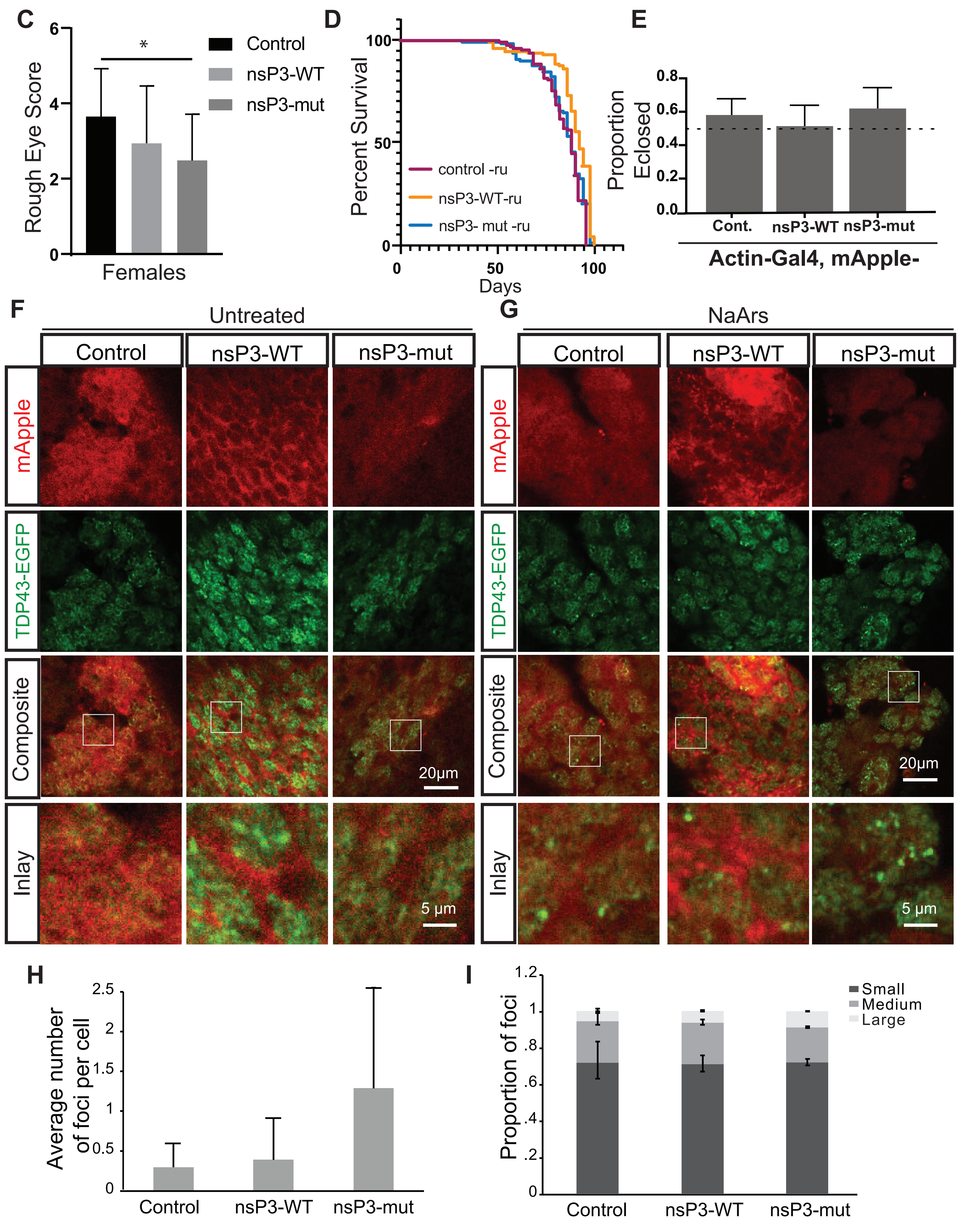

C
Rough Eye Score
Females
Control
nsP3-WT
nsP3-mut
D
Percent Survival
Days
control -ru
nsP3-WT-ru
nsP3- mut -ru
E
Proportion Eclosed
Cont.
nsP3-WT
nsP3-mut
Actin-Gal4, mApple-
F
Untreated
G
NaArs
Control
nsP3-WT
nsP3-mut
mApple
TDP43-EGFP
Composite
Inlay
20μm
5 μm
H
Average number of foci per cell
Control
nsP3-WT
nsP3-mut
I
Proportion of foci
Small
Medium
Large
Control
nsP3-WT
nsP3-mut

**Supplemental Figure 3: nsP3 prevents SG formation in larval Drosophila brains.** A) Representative images of 3rd instar larval brains expressing rin-sfGFP, and Actin-Gal4, nsP3-mut either untreated, or 1hr at 42C or 2hrs in 500uM NaArs. B) Representative images of 3rd instar larval brains expressing rin-sfGFP, Actin-Gal4 and either mApple control, nsP3-WT or nsP3-mut in the presence of 10uM thapsigargin (TG) for 90 minutes. C) Quantification of rough eye phenotypes of female flies expressing GMR-Gal4; TDP43-EGFP and either control, nsP3-WT, or nsP3-mut (Control n= 24, nsP3-WT n= 22, nsP3-mut n= 22). One-way ANOVA and Tukey's multiple comparisons test. *$p<0.05$. D) Survival assay of Tub5-GS, mApple control, nsP3-WT, or nsP3-mut flies without RU-486 at 24C. (control n= 91, nsP3-WT n= 89, nsP3-mut n= 92). Log-rank Mantel-Cox test. E) Bar graph represents proportion of eclosed flies from Actin-Gal4/Cyo crossed to homozygous mApple control, nsP3-WT, or nsP3-mut flies that were mApple+, +/- 95% C.I.s. Chi Square Analysis, (control n=101, nsP3-WT n= 66, nsP3-mut n=58). Dotted line is expected assuming no synthetic lethality caused by gene expression. F-G) Representative images of 3rd instar larval brains expressing Asense-Gal4; TDP43-EGFP, and mApple control, nsP3-WT, or nsP3-mut, incubated for 2 hrs untreated (F) or in 500uM NaArs (G). H) Quantification of average number of foci/cell/image (n=images: control n= 10, nsP3-WT n=14, nsP3-mut n=12) in (G). Bars represent mean +/- standard deviation. Multiple T-tests, no significance. I) Quantification of foci size distribution from images in (G). Bars represent proportion of total foci +/- 95% confidence intervals. Chi-square analysis, (control n= 71, nsP3-WT n=221, nsP3-mut n=479), not significant.

Supplemental Figure 4

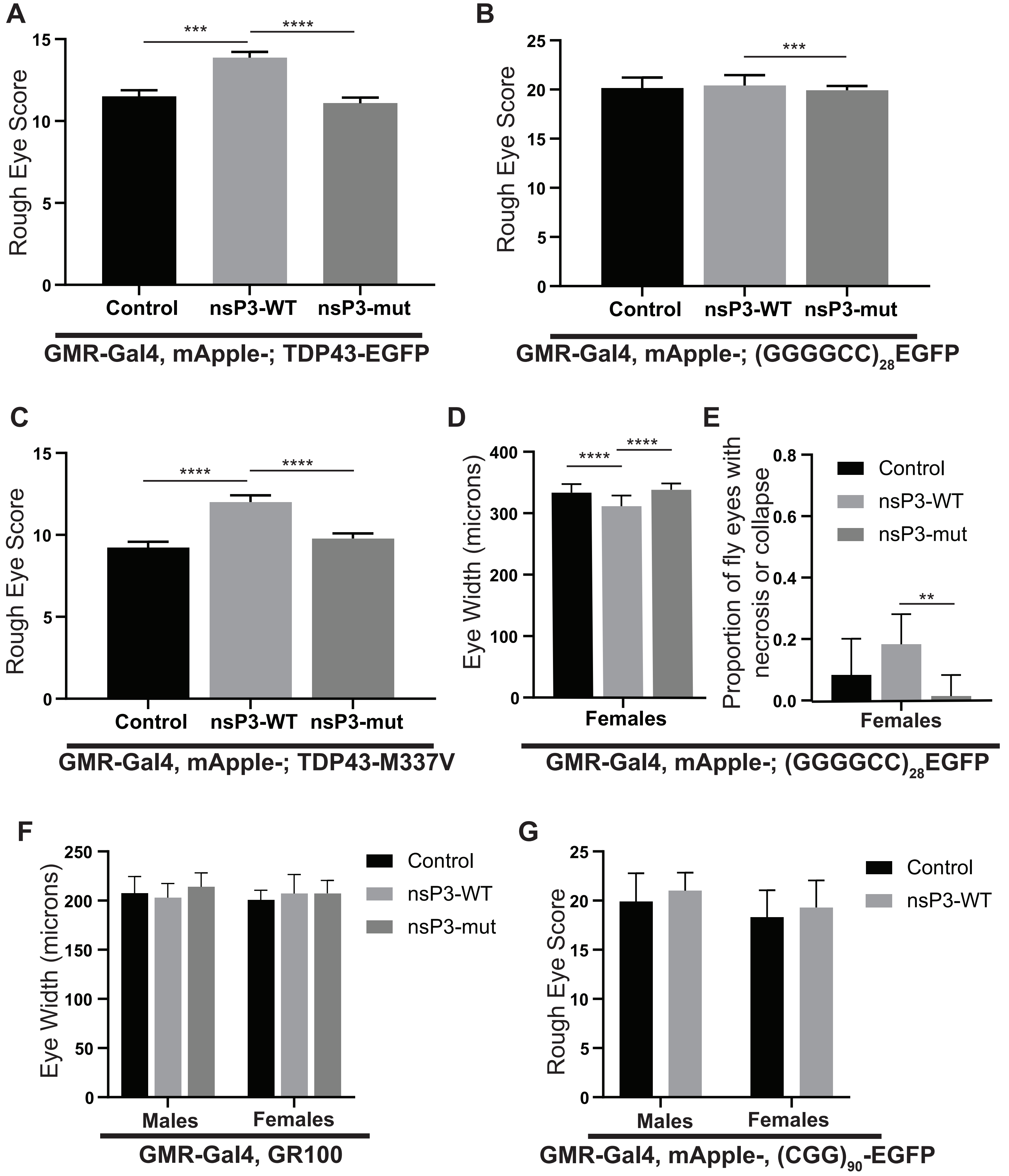

**Supplemental Figure 4: nsP3 enhances toxicity in ALS fly models.** A-C) Quantification of female GMR-Gal4/ mApple control, nsP3-WT, or nsP3-mut + A) TDP43TDP43-EGFP (control n= 60, nsP3-WT n= 52, nsP3-mut n= 88) ****p<0.0001, B) (GGGGCC)28-EGFP (control n=48, nsP3-WT n=82, nsP3-Mut=71) **p<0.01, and C) TDP43-M337V (control n= 37, nsP3-WT n= 38, nsP3-mut n= 54) ****p<0.0001 rough eye phenotype. A-C) One-way ANOVA with Tukey's multiple comparison tests. Quantification of (GGGGCC)28-EGFP female D) eye width (control n= 23, nsP3-WT n= 22, nsP3-mut n= 25) ****p<0.0001, and E) proportion of necrosis positive female eyes (control n=48, nsP3-WT n=82, nsP3-Mut=71). F) Quantification of GR100 eye width (control male n= 24, female n= 12; nsp3-WT male n= 17, female n= 10; nsP3-mut male n= 11, female n= 11). D & F: Two-way ANOVA (see Figure 6F), and Tukey's multiple comparison tests. E: Fischer's exact test with Bonferonni correction for multiple comparisons. ***p<0.001. G) Quantification of GMR-Gal4, (CGG)90-EGFP, mApple-x rough eye phenotype (males control n= 45, nsP3-WT n= 14; females control n= 41, nsP3-WT= 23). Two-way ANOVA with Sidak's multiple comparisons test.

Supplemental Figure 5

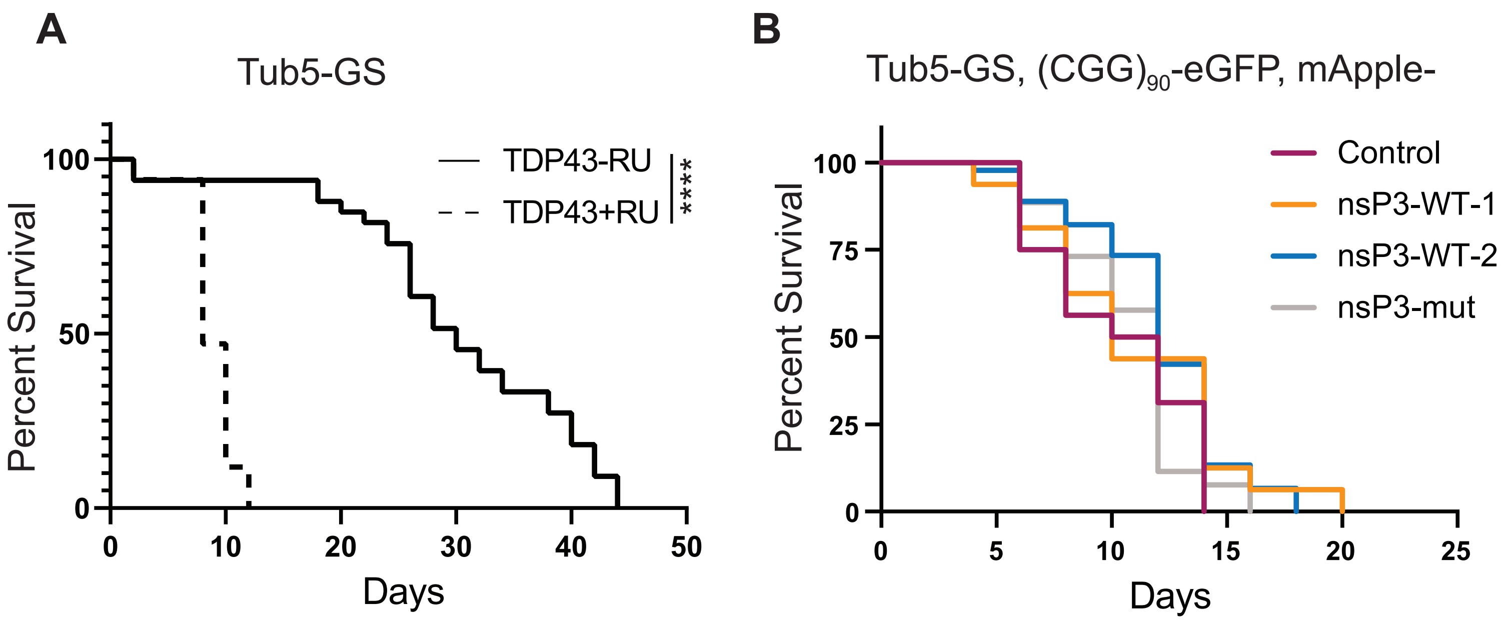


**Supplemental Figure 5. nsP3 has no effect on survival of flies expressing CGG repeats.** A) Survival assay of Tub5-GS, TDP43-GFP +/- RU B) Survival assay of Tub5-GS, (CGG)90-EGFP, crossed to mApple control, nsP3-WT, or nsP3-mut. (control n= 16, nsP3-WT-1 n=16, nsP3-WT-2 n=45, nsP3-mut n=26). Log-rank Mantel-Cox test with Bonferroni corrections for multiple comparisons.
